# Measurements of 3-D morphology of individual red blood cells upon localized photothermal effects using gold nanorods

**DOI:** 10.1101/2019.12.11.872218

**Authors:** Chanwoo Song, Gu-Haeng Lee, JunTaek Oh, Moosung Lee, Seungwoo Shin, SangYun Lee, Yoonkey Nam, YongKeun Park

## Abstract

We investigated the *in-situ* photothermal response of human red blood cells (RBCs) by combining photothermal heat generation and 3D quantitative phase imaging techniques. Gold-nanorod-coated substrates were excited using near-infrared light to generate local heat to RBCs, and response was measured by imaging 3-D refractive index tomograms of cells under various near infrared (NIR) excitation conditions. On photothermal treatment, cell morphology changed from discoid to crescent shapes, cell volume and dry mass decreased, and hemoglobin concentration increased. We also investigated the irreversible deformation of RBCs when multiple intense excitation shocks are applied. These results provide a new understanding of thermodynamic aspects of cell biology and hematology.

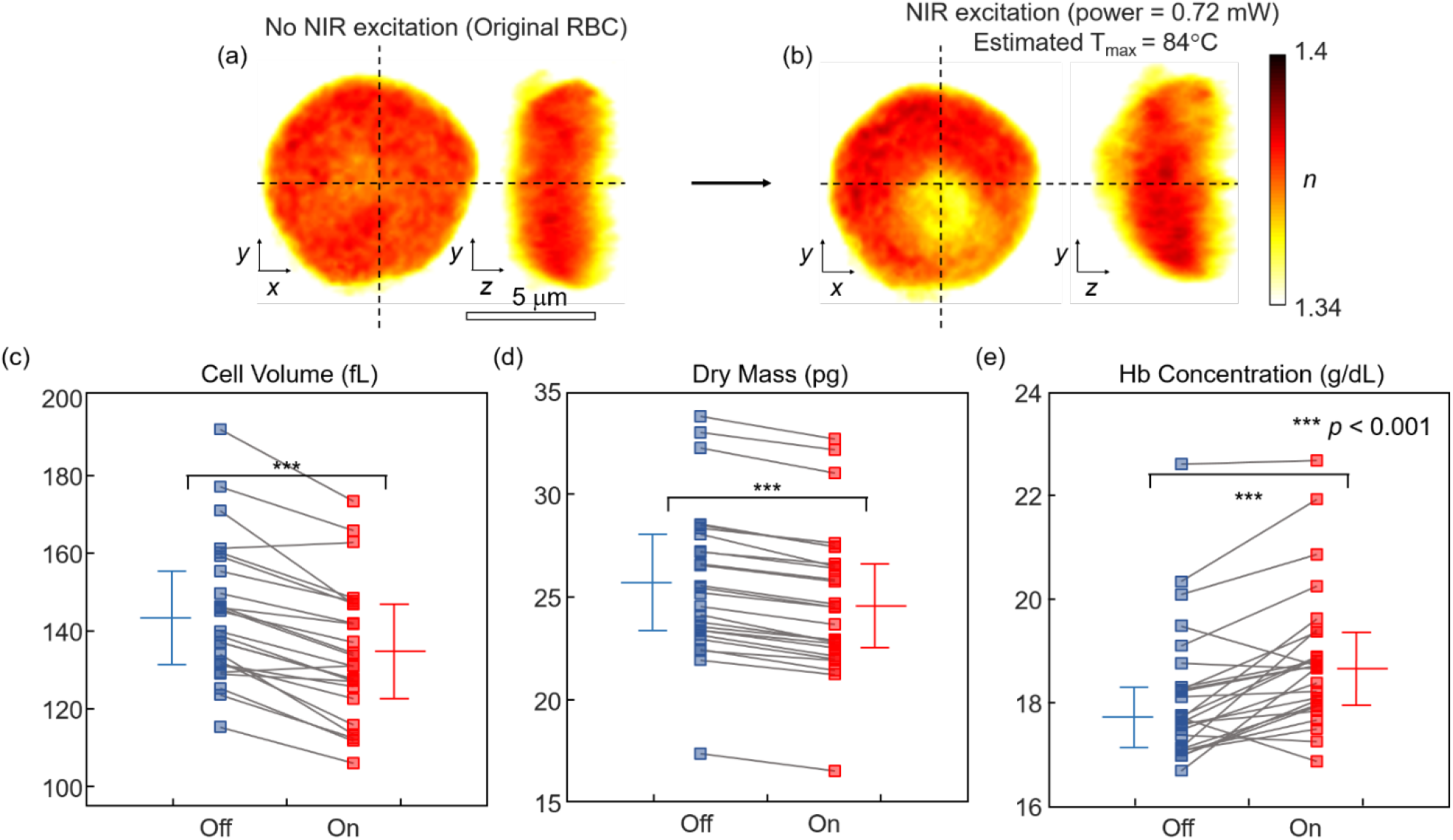

## INTRODUCTION

Alterations in the biophysical properties of red blood cells (RBCs) are directly related to various diseases such as sickle cell disease^1,2^, malaria ^3–6^, and diabetes ^7,8^, and can be potential indicators for diagnosing various diseases ^9,10^. The thermal response of RBCs can be an effective indicator for the pathophysiology of various diseases. Even though it is well known that the pathophysiology of many RBC related diseases is temperature related ^3^, most previous studies have been limited to a small temperature range (between room temperature and body or fever temperature) ^3^. Furthermore, a systematic investigation of the cellular response to local heating of individual cells, particularly using three-dimensional imaging, has not been explored. This is unfortunate because it could provide valuable information to connect the pathophysiology of diseases to thermodynamics, and unveil complex, non-equilibrium thermodynamic behavior ^11–15^.

The study of the thermal response of RBCs at an individual cell level has been hindered by technical limitations of heat treatment and imaging methods. First, local heat treatment of biological cells is inaccessible using a conventional heating chamber. Most previous thermodynamic experiments on RBCs have relied upon ensemble averaging of multiple cells, such as a case using a microfluidic pipette ^16^. Furthermore, rapid three-dimensional (3-D) time-lapse imaging of RBCs is required to study cellular response upon local heating. However, imaging the 3-D dynamics of RBCs over a long time at a high frame rate is challenging for either conventional bright-field or fluorescence-based microscopy. Owing to these limitations, conventional imaging approaches have provided limited access to the morphological and biochemical properties of RBCs, such as lateral shapes and out-of-membrane fluctuations ^17–19^.

To directly investigate the in-situ thermal response of individual RBCs, we present a novel combination of two optical approaches: photothermal heat generation using gold nanorods (GNRs) and 3-D rapid quantitative imaging using a 3-D quantitative phase imaging technique (QPI). The GNR-coated coverslips, on which RBCs are located, are locally excited by illuminating a tightly focused optical spot using an 808 nm wavelength near-infrared (NIR) laser. The NIR excitation beam locally generates heating of GNRs via surface plasmonic resonance ^20,21^. Due to conduction, the temperature of the RBCs rapidly increases and induces a thermal response from individual RBCs. Then, in order to measure the 3-D response of individual RBCs, we employ optical diffraction tomography (ODT), one of several 3-D QPI techniques ^22,23^. QPI is a label-free imaging technique using the refractive index (RI) for intrinsic and quantitative imaging contrast ^22,24^. By utilizing interferometric or non-interferometric imaging systems, QPI has been applied to various applications, including hematology ^25–28^, malaria infection ^3,29,30^. neuroscience ^31,32^, cell biology ^33–35,36^, reproductive biology ^37^, diabetes ^7^, and biotechnology ^38^, due to its label-free, live, and rapid imaging capability.

Here, by measuring the 3-D RI distribution of an RBC being photothermally excited, various biophysical parameters of the target cell are systematically investigated. We observed and quantitatively analyzed the morphological and biophysical changes of individual RBCs under local heat exposure *in situ*. In this report, we suggest that the integration of the ODT technique and photothermal heat generation serves as a novel method for extensive study of various single-cell thermodynamics.

## METHODS AND MATERIALS

### Experimental Setup

The experimental setup and the analysis procedure are illustrated in Fig. 1. RBCs dispersed in Alsever’s solution were sandwiched between two No.1-thickness coverslips (Fig. 1(a)). The bottom coverslip was coated with cylindrically symmetric ellipsoidal GNRs, whose diameters were 14 nm and 50 nm on the minor and major axes, respectively (Fig. 1(b)). The GNRs were designed and optimized to generate heat diffusion under NIR light at 808 nm and effectively raise the temperature of a sample due to its high absorption and heat conversion efficiency ^39^ (Fig. 1(c)).

**Fig. 1.**
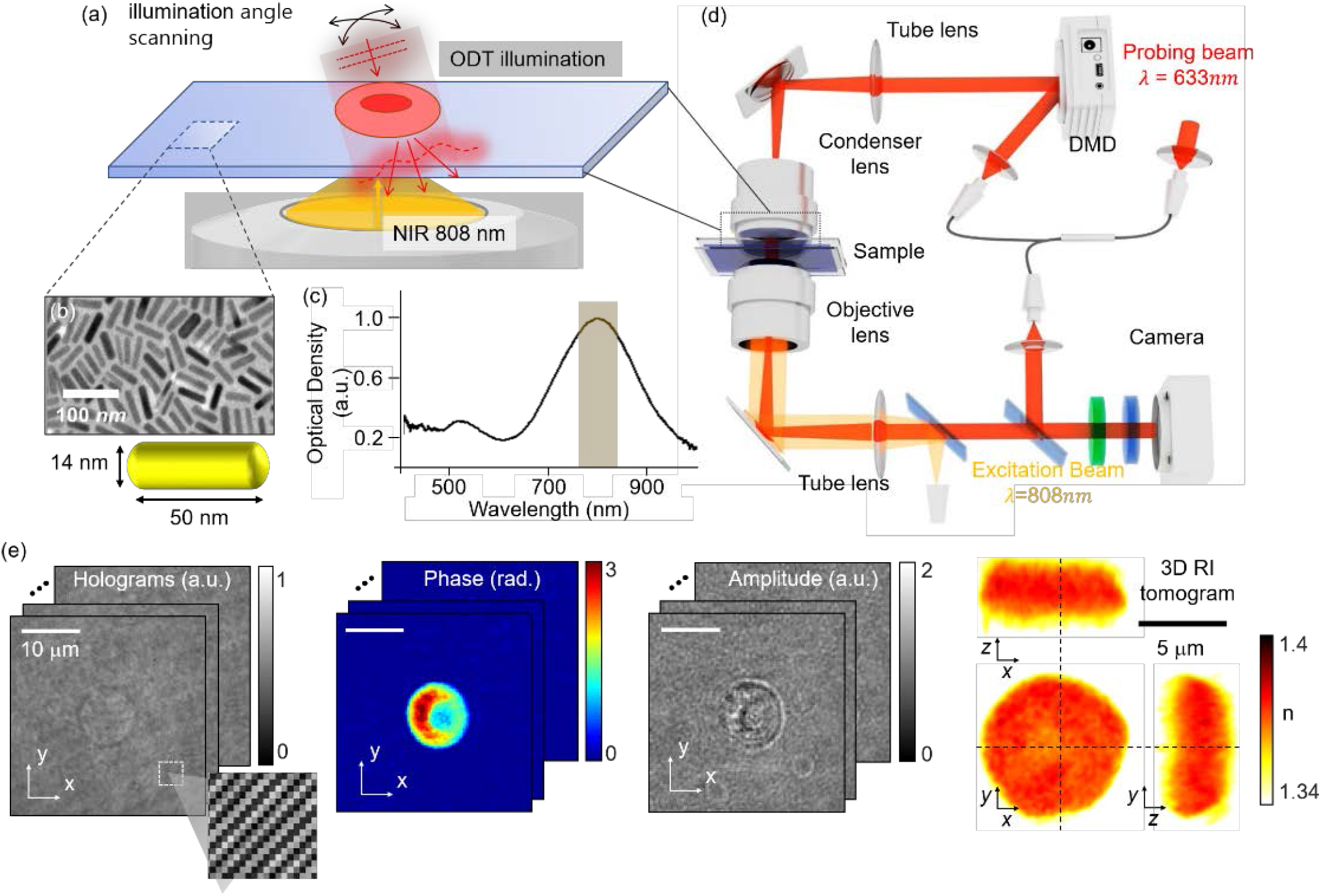
(a) Experimental schematics. RBC samples were heated by GNRs excited by NIR illumination. (b) Transmission electron microscopy of GNRs and the GNR shape. (c) Measured absorption spectrum of the GNR coated coverslip^40^. (d) Optical setup. The 3-D morphology of individual RBCs is measured using ODT equipped with a DMD, while the sample is excited by the NIR laser. (e) ODT reconstruction process. The 3-D RI tomogram is reconstructed from the measured multiple 2D holograms of a sample.

In order to simultaneously increase the local temperature and probe the 3-D response of the heated RBC, we used an ODT system (Fig. 1(d)) combined with an NIR laser for photothermal excitation and a visible laser for imaging ^39^. To raise the temperature of individual RBCs, an NIR laser diode (*λ* = 808 nm, M9-808-015, Thorlabs Inc., USA) was used to excite GNRs. The NIR laser beam was focused at the center of a target RBC, with a beam diameter of 8.5 *μ*m. The temperature increase was calibrated using data obtained with the GNR-coated coverslip in the absence of cells ^40^. We assumed that the temperature of target RBCs increases immediately upon NIR illumination, based on the nature of the rapid thermal equilibrium of a micro-system ^41,42^.

To measure 3-D images of individual RBCs upon thermal treatment, an ODT system equipped with a DMD was utilized. A He-Ne laser (*λ* = 633 nm, 15 mW, Thorlabs Inc., USA) was split into a reference beam and a sample beam using a 2×2 single-mode optical fiber coupler. To control illumination angles of the sample beam, a DMD (DLP LightCrafter 6500, Texas Instruments Inc., USA) was located at the conjugate plane to a sample ^43^. Time-multiplexed hologram patterns were projected onto the DMD, to control the illumination angle of the beam impinging onto a sample, with high speed (100 Hz) and accuracy ^41,42^. The modulated illumination beam was projected onto a sample using a tube lens (TL1, *f* = 250 mm) and a water-immersion condenser lens (CL, NA = 1.1 LUMPLN, 60×, Olympus Inc., Japan). To collect the diffracted beam from the sample, the water-immersion objective lens (OL, NA = 1.2 UPLSAPO, 60×, Olympus Inc., Japan) and a tube lens (TL2, *f* = 180 mm) were used. The sample beam was then imaged on to a camera plane, where it interfered with a reference beam with a slight tilt, resulting in the formation of a spatially modulated interference pattern. The interference patterns were recorded using a CMOS sensor (FL3-U3-13Y3M-C, FLIR Systems Inc., USA). To avoid the detection of the NIR excitation laser beam at the camera, a short pass filter (edge wavelength = 750 nm, Thorlabs Inc., USA) was placed in front of the camera.

From the recorded interference patterns of a sample illuminated at various incident angles, a 3-D RI tomogram was reconstructed using ODT principles (Fig. 1(e)) ^44^. First, optical field images, consisting of both the amplitude and phase images, were retrieved from the interference patterns using a phase retrieval algorithm ^45,46^. Then, retrieved optical field images were mapped into a 3-D Fourier space, based on the Fourier diffraction theorem and the Rytov approximation ^23,44^. Due to the limited numerical apertures of both the condenser and objective lenses, side scattering information was not collected, resulting in a missing cone problem. To remedy the missing cone problem, the Gerchberg-Papoulis (GP) algorithm based on a non-negativity constraint was applied ^47,48^. The iteration number was set to 100, and it was assumed that the sample RI is higher than the medium RI. The theoretical lateral and axial resolutions are 110 nm and 360 nm, respectively ^49^. The details of ODT with the Matlab reconstruction code and the calibration of local heating can be found elsewhere ^40,50–52^.

### Synthesis of gold nanorods

A seed-mediated method was utilized to synthesize GNRs ^53^. At room temperature, 2.5 ml of 0.2 M cetyltrimethylammonium bromide (CTAB, Sigma), 2.5 ml of 0.5 mM HAuCl4 (Sigma), and 300 μL of ice-cold 0.01 M NaBH4 (Sigma) were mixed in an ultrasonication bath and used as a seed solution. After aging the seed solution for 2 h, seeds were grown to a rod shape in growth solution for 30 min at room temperature. The growth solution was a mixture of 5 ml 0.2 M CTAB, 5 ml of 1 mM HAuCl4, 250 μL of 4 mM AgNO3 (Sigma), 70 μL of 78.84 mM ascorbic acid (Sigma) and 12μL of seed solution. The GNRs were then concentrated by centrifugation at 10,000 RPM (10,200 RCF) and resuspended in ultrapure water in order to remove the surfactant. The GNRs were coated with polyethylene glycol (mPEG-SH, MW 5000, Nanocs) with a ratio of 3 optical density (O.D.) of GNR to 3 mg/ml of PEG in water solution for 12 h at room temperature. Using a dialysis kit (Thermo Scientific), excess PEG was removed for two days. We measured the zeta potential of PEG-coated GNR by Zetasizer Nano ZS (Malvern).

To ensure a clean surface, glass substrates were cleaned with acetone, isopropyl alcohol, and deionized water for 5 min each with ultrasonication. Layer-by-layer coating was performed on each substrate to create a positively charged surface. For the substrate treatment, we prepared 10 mg/ml of poly(sodium 4-styrenesulfonate) (PSS, MW~70,000, Aldrich) and poly(allylamine hydrochloride) (PAH, MW~17,500, Aldrich) in 10 mM NaCl solution. Then, the glass substrates were treated with PSS and PAH solution for 3 cycles of 5 min each and finalized with PAH. A 1 O.D. GNR solution (0.2 ml/cm^2^) with a zeta potential of −37.5 mV was dispersed on the substrates for 12 h. Slight mismatch of absorption peak was tuned by additional layer-by-layer coating on GNR coated substrates. The average length and diameter of synthesized GNRs are 50 nm and 14 nm, respectively (aspect ratio of 3.60).

We measured the absorbance spectrum and the absorbance value at 808 nm by vis-NIR spectroscopy (Ocean Optics). The absorption spectrum of the GNRs was observed to have a maximum peak at around 808 nm (Fig. 1(c)). Using absorbance microscopy, the optical properties of the sample were also measured. The average absorbance of the samples upon excitation was shown to have an O.D. of 0.127 O.D. and an absorbance ratio of 25.4%^40^.

### Preparation of red blood cell samples

Blood was extracted using a lancet. A total of 5 μL of blood was collected from healthy volunteers using a pipette, and then diluted in 1 mL of Alsever’s solution (A3551, Sigma-Aldrich, Missouri, United States). For each measurement, 50 μL of the diluted solution was dropped onto a GNR-coated coverslip and covered with another bare coverslip (24×40 mm, C024501, Matsunami, Ltd, Japan). The sample was loaded on a stage of the optical setup and measured. All the experimental protocols were approved by the institutional review board of KAIST (IRB project: KH2015-37).

### Morphology analysis of individual RBCs

The morphological and biophysical parameters of individual RBCs were retrieved from measured 3-D RI tomograms. Because the RI contrast of RBCs provides their hemoglobin concentration distribution ^54^, ODT enables label-free quantification of RBC parameters in a non-invasive manner ^55,56^. The following cell parameters were retrieved: cell volume, hemoglobin (Hb) concentration, and cellular dry mass.

The cell volume was calculated from the number of voxels with an RI value higher than a threshold value of 1.350. The Hb concentration of each cell, [Hb], was calculated from the measured RI values of the RBC cytoplasm. Due to a linear relationship between the mean RI value of non-aqueous cytoplasmic contents and its mean concentration ^57,58^, the Hb concentration of an RBC can be calculated from the mean RI contrast 〈Δ*n*〉:

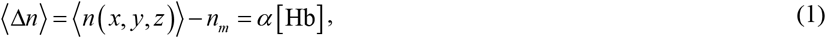

where *n_m_* is the RI of a medium, *α* is the refraction increment, a coefficient between the relationship between the mean RI of RBC cytoplasm and Hb concentration. In this work we used a value of 0.18 mL/g for *α* ^54,59^. The dry mass of individual RBCs, the mass of non-aqueous cytoplasmic contents, can also be obtained from integration of the Hb concentration over a cytoplasmic volume:

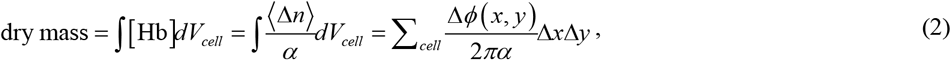

where Δ*ϕ*(*x, y*) is a measured phase delay in a given coordinate and Δ*x*Δ*y* is the area in the field of view corresponding to the pixel area of the camera.

### Statistical Analysis

For a statistical comparison of the mean RBC parameters of the original and heat-treated RBC groups, a Wilcoxon rank-sum test was utilized. In experiments based on individual cell tracking, the paired Wilcoxon rank-sum test was employed to address continuous alterations in the measured parameters of individual RBCs under the heat treatment process. Throughout the manuscript, all the retrieved RBC parameters are given in the form mean ± std, and a *p*-value of less than 0.05 was regarded as statistically significant.

## Results

### Effects of localized heating on RBCs

To demonstrate the effects of localized heating on RBCs, a 3-D RI tomogram of an individual RBC was measured before and two seconds after NIR excitation (Fig. 2). Laser was continuously irradiated during image acquisition, which requires 0.3 s. Representative images before and after NIR excitation are shown in Figs. 2(a-b). Before excitation, cells exhibit a characteristic biconcave shape (Fig. 2(a)). After NIR excitation (0.72 mW), a significant morphological alteration causes the overall shape to be crescent.

**Fig. 2.**
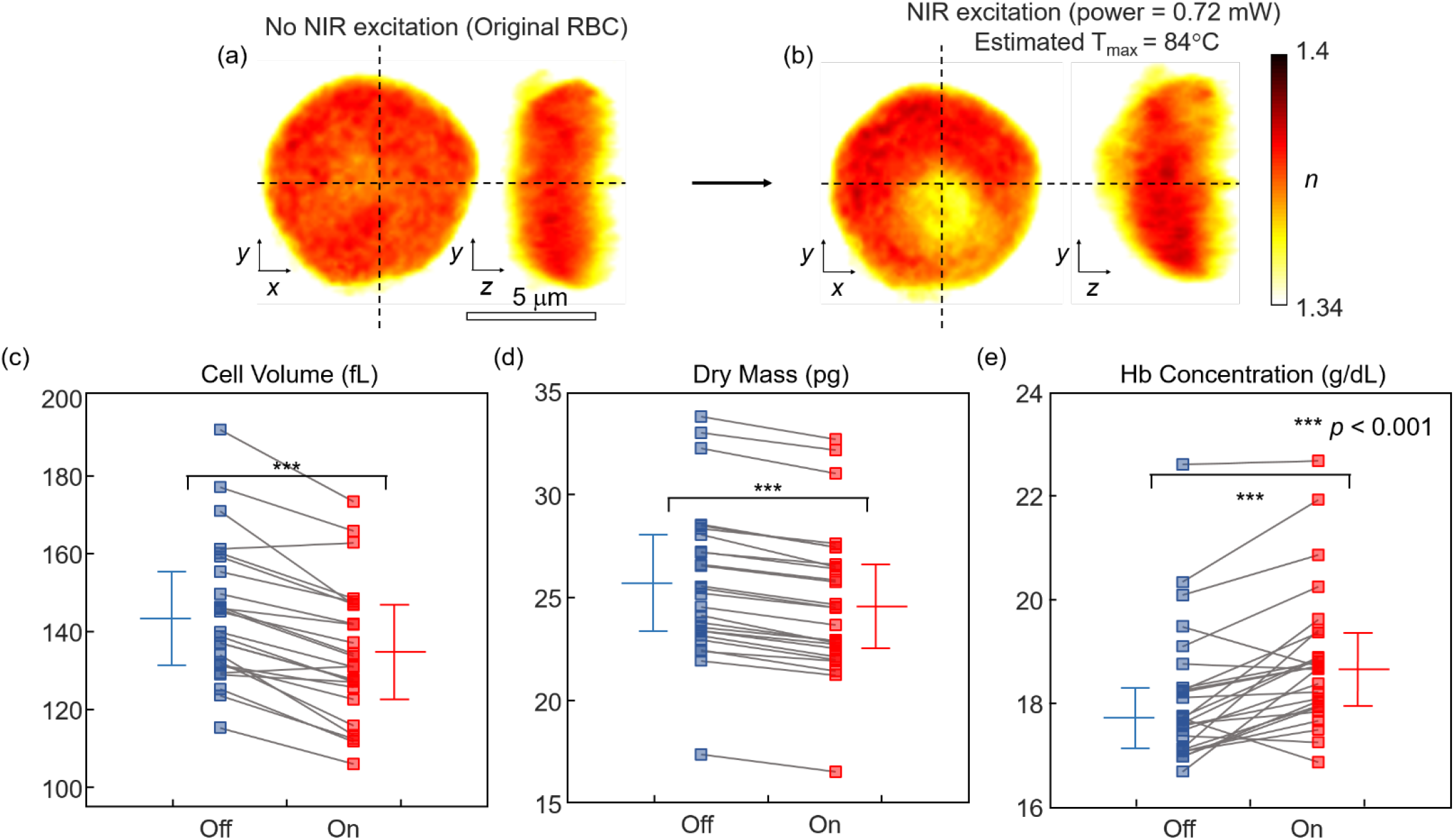
(a,b) Reconstructed 3-D RI tomograms of representative RBCs before and after NIR excitation. (c) Cell volume, (d) dry mass, (e) Hb concentration for all measured RBCs before and after NIR excitation exposure with a power of 0.72mW (*n* = 33). Each square denotes an individual RBC measurement. The horizontal lines indicate the mean values, and error bars represent standard deviation.

For a clearer comparison, these morphological and biochemical RBC responses were quantified and statistically analyzed (Figs. 2(c-e)). In particular, we retrieved the statistics of volume, dry mass, and Hb concentration of 33 RBCs from their measured 3-D RI tomograms. The mean volume of RBCs significantly decreased from 142.41 ± 20.53 fL at room temperature (25 °C) to 132.35 ± 19.81 fL after the excitation from room temperature (*p* < 0.001) (Fig. 2(c)). The total dry mass also decreased significantly from 25.73 ± 3.67 pg to 24.84 ± 3.61 pg after the excitation (*p* < 0.001) (Fig. 2(d)). These data indicate that the localized photothermal heat generation induced the efflux of internal RBC components. To determine whether the major effused component is water or hemoglobin, we compared the statistics of the mean Hb concentration of RBCs before and after NIR excitation (Fig. 2(e)). The retrieved mean Hb concentrations of RBCs at RT and after localized photothermal treatment are 18.12 ± 1.31 g/dL and 18.83 ± 1.35 g/dL, respectively, indicating a significant difference (*p* < 0.001). From these data, we confirmed that about 70% of volume decrease is contributed by water.

### Deviation in RBC parameters is directly related to power of excitation beam for localized heat treatment

To determine whether there is a significant correlation between the amount of photothermally generated heat and the RBC physiological changes, we examined the morphological and biochemical parameters of RBCs as a function of NIR illumination power (Fig. 3). We controlled the NIR laser power from 0.48 to 0.84 mW with an interval of 0.12 mW, and quantified the relative parameter changes before and after the photothermal treatments. Same imaging procedure with what mentioned at section 3.1 was adopted in this experiment.

**Fig. 3.**
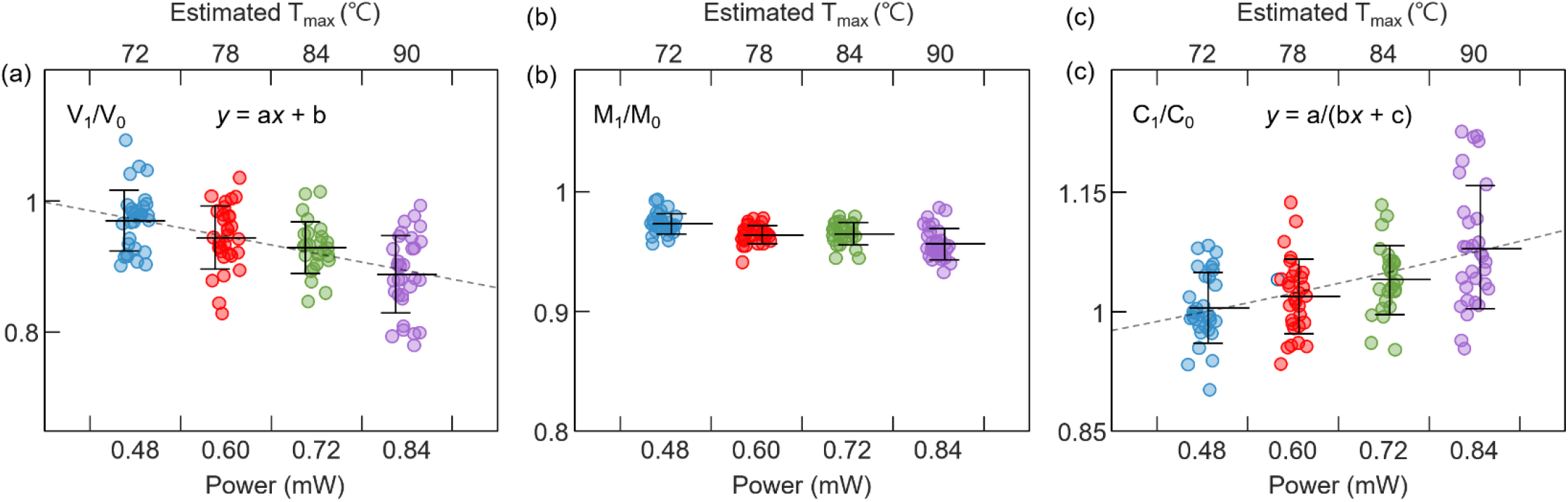
(a) Relative RBC volume before and after heat treatment (V_1_/V_0_) for each RBC with respect to NIR excitation power. Gray dotted line is linearly fitted graph that describe relationship between average value of V_1_/V_0_ and the excitation power. Coefficients are statistically defined as a = −21.7833 [mW^−1^] and b = 1.286 with p-values of 0.0167 and 1.30×10^−3^ for each parameter, respectively. (b) Relative RBC dry mass before and after heat treatment (M_1_/M_0_) for each RBC with respect to NIR excitation power. (c) Relative RBC Hb concentration before and after heat treatment (C1/C0) for each RBC with respect to the NIR excitation power. Gray dotted line is rationally fitted graph that describe relationship betwwen average value of C1/C0 and the excitation power. Coefficients are statistically defined as a = 0.0154 [mW^−1^], b = −0.30209 and c = 82.453 with p-values of 5.70×10^−4^, 0.0164, and 2.11×10^−4^, respectively. Each circle denotes an individual RBC measurement; n = [33, 30, 33, 33] for [0.48, 0.60, 0.72, 0.84] mW NIR power respectively. The center horizontal line indicates the mean values, and the error bar indicates the standard deviation.

As shown in Fig. 3(a), we found a significant negative correlation between NIR laser power and volume. The volume decrease ratio per power was −21.78 /mW below 0.84 mW. In contrast, the dry mass change with power level was inconsistent (Fig. 3(b)). Based on these statistics, the power dependence of the hemoglobin concentration was rationally fitted [Fig. 3(c)]. As expected, the fitted curve showed significant positive, rational relationship (R^2^ > 0.99). Overall, these trends agree well with the single-cell statistics in Fig. 2, and indicate that volume decrease and Hb concentration increase are significant features of RBCs subjected to heat treatment.

### Reversibility of the RBC alteration due to localized heat treatment

We further examined the reversibility of the photothermal response of individual RBCs (Fig. 4). We illuminated a 2.3-second pulse train NIR laser beam with a time interval of 4.6 s, and analyzed the response of individual RBCs (Fig. 4(a)). Same imaging procedure from section 3.1 was adopted for each steps of NIR exposure.

**Fig. 4.**
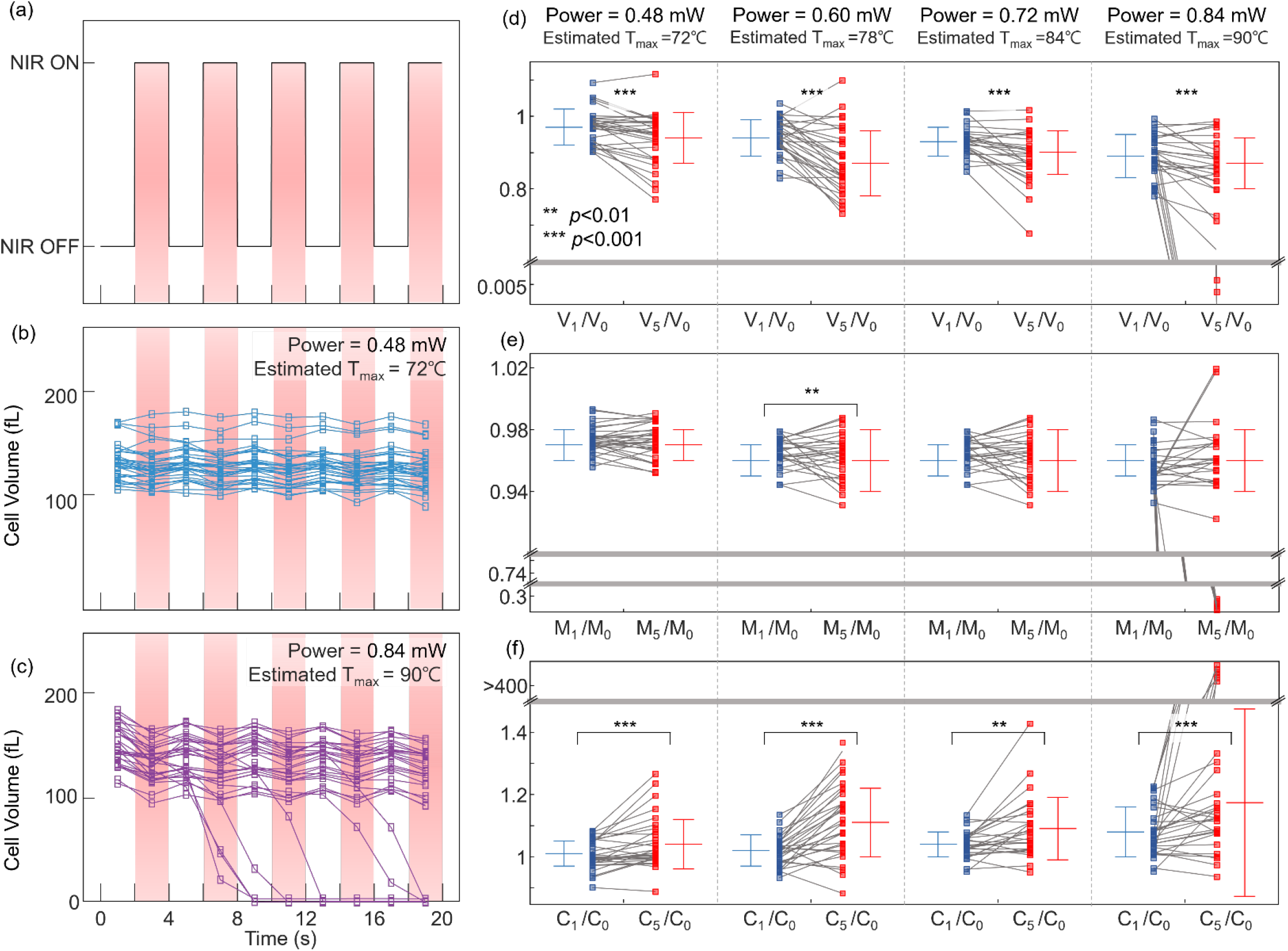
(a) NIR excitation power profile with respect to time is presented. Individually traced plot of cell volume of RBCs with NIR power of (b) 0.48 mW and (c) 0.84 mW. Calculated (d) cell volume (V_*i*_V_0_), (e) dry mass (M_*i*_/M_0_), and (f) protein concentration (C_*i*_/C_0_), where *i* indicates the *i*th step of heat exposure. Each blue and red square denotes an individual RBC measurement for the first heat exposure step and the fifth heat exposure step. The center horizontal line indicates the mean values, and the error bar indicates the standard deviation; *n* = [33, 30, 33, 33] for [0.48, 0.60, 0.72, 0.84] mW of NIR power respectively. ** and *** denote that the *p*-value is smaller than 0.01 and 0.001, respectively.

Under low NIR power (0.48 mW, estimated T_max_ = 72°C), we observed that all the RBCs exhibited reversible volume decreases after 5 cycles of heat treatment (Fig. 4(b)). Decreased volume of the RBCs due to heat addition was recovered instantaneously when the excitation laser was turned off (Fig. 4(b), Supplementary Video 1). However, upon high power NIR illumination (0.84 mW, estimated T_max_ = 90°C), an irreversible recovery was observed after multiple heat exposure (Fig. 4(c), Supplementary Video 2). Notably, we observed that several RBCs burst when exposed to the 0.84 mW NIR beam (Supplementary Video 3).

We studied the heat-dependent irreversibility of RBCs by comparing the relative parameter ratios after the first and fifth exposures (Figs. 4(d-f)). The relative volume ratio statistics indicate dramatic irreversible volume decreases after five times of NIR excitation (Fig. 4(d)). This may be related to the fact that the estimated maximum temperature exceeded the typical range of protein denaturation temperature, around 40°C. In contrast, the dry mass ratio remained almost constant except when RBCs burst at high heat levels (Fig. 4(e)). As a result, the irreversible concentration increase was more dramatic with stronger NIR laser power (Fig. 4(f)). Overall, our results suggest that irreversible changes in RBCs due to heat treatment are correlated with volumetric decrease and concentration increase.

### Discussions and Conclusion

Here we presented *in situ* observations of morphological and biochemical alterations of RBCs subjected to localized photothermal treatment. Localized photothermal treatment was applied to individual RBCs by using NIR excitation of GNRs. The corresponding alterations of individual RBCs were precisely and quantitatively measured in 3-D using ODT. Our study revealed significant morphological changes in RBCs after localized photothermal treatments, including volume decrease, dry mass loss, and concentration increase. We also investigated the reversibility of the RBC physiological changes driven by photothermal excitation. We observed three key quantitative parameter changes depending upon local temperature increment: decreasing volume, decreasing dry mass, and increasing protein concentration. Although an overall understanding of thermally altering RBC properties remains a great challenge, our approach and findings provide an important step to address biophysical models for RBC thermodynamics. These methods can also be utilized in the context of cellular physiology.

The physiological origin of these changes is still unclear, so we anticipate that the following hypotheses can be tested in future studies. First, the volume decrease of RBCs under heat treatment may be explained in association with larger water permeability across the RBC membrane at higher temperature ^60,61^. Second, the dry mass loss of RBCs at temperatures higher than 40°C is very likely a result of protein denaturation in the RBCs. In order to validate these hypotheses, biochemical analysis tools should be used for further exploration.

Although we focused on the heat response of RBCs, the proposed method is generally applicable to studying the physiology of other types of cells. Because GNRs have been conventionally applied in photothermal therapy, it would be intriguing to investigate single-cell studies related to cancer treatment ^20^, neuron signal modulation ^56,62,63^, and neuron growth ^64^. Furthermore, the responses of individual RBCs at high temperatures can also be utilized for in vivo photothermal therapy ^65^. Also, the throughput of the proposed system can be extended by combining a wide-field imaging platform. In such systems, cell-cell interactions under local heat treatment could also be investigated. Overall, the proposed method opens new horizons for understanding the thermodynamic physiology of biological systems.

## Acknowledgments

This work was supported by KAIST, BK21+ program, Tomocube, and National Research Foundation of Korea (2017M3C1A3013923, 2015R1A3A2066550, 2018K000396). The authors thank Dr. Weisun Park for helpful discussions.

